# Trade-offs between baseline thermal tolerance and thermal tolerance plasticity are much less common than it appears

**DOI:** 10.1101/2022.11.30.518595

**Authors:** Alex R. Gunderson

## Abstract

Thermal tolerance plasticity is a core mechanism by which organisms can mitigate the effects of climate change. As a result, there is a need to understand how variation in tolerance plasticity arises. The Baseline Tolerance/Plasticity Trade-off Hypothesis (hereafter referred to as the Trade-off Hypothesis, TOH) has recently emerged as a potentially powerful explanation. The TOH posits that organisms with high baseline thermal tolerance have reduced thermal tolerance plasticity relative to those with low baseline tolerance. Many studies have found support for the TOH. However, this support must be regarded cautiously because the most common means of testing the TOH can yield spurious ‘trade-offs’ due to regression to the mean. I acquired data for 25 previously-published analyses that supported the TOH at the intraspecific level and reanalyzed them after applying a method that adjusts plasticity estimates for regression to the mean. Only six of the 25 analyses remained statistically significant after adjustment, and effect size and variance explained decreased in all cases. The few data sets in which support for the TOH was maintained after adjustment point to areas of future study, but are too few to make generalizations at this point. In sum, regression to the mean has led to a substantial overestimation of support for the TOH and must be accounted for in future tests of the hypothesis.

## Introduction

The capacity to physiologically adjust to temperature change through phenotypic plasticity is thought to influence vulnerability to anthropogenic warming (Gunderson, Dillon, & Stillman, 2017; Huey et al., 2012; Morley, Peck, Sunday, Heiser, & Bates, 2019; Seebacher, White, & Franklin, 2015; Somero, 2010). Uncovering how and why taxa differ in physiological plasticity is therefore of great interest (Kelly, 2019; Pottier, Burke, Drobniak, Lagisz, & Nakagawa, 2021; Rohr et al., 2018). Recently, the Trade-off Hypothesis (TOH) has emerged as a potentially powerful mechanism to explain variation in the plasticity of thermal tolerance limits (Stillman, 2003; van Heerwaarden & Kellermann, 2020). This hypothesis posits that taxa with high baseline levels of thermal tolerance will exhibit low levels of thermal tolerance plasticity. From a mechanistic perspective, the TOH is usually discussed in terms of phenotypic constraint. Thermal limits can only be so high, and therefore plasticity must be reduced as baseline tolerance rises. The implications of this hypothesis, if and when true, are many, and touch not just on global change biology but fundamental aspects of physiological ecology and evolution (reviewed in van Heerwaarden & Kellermann, 2020).

Support for the TOH has been found in several studies at both the intraspecific and interspecific levels (Barley et al., 2021; Comte & Olden, 2017; van Heerwaarden & Kellermann, 2020; see also Table 1). Nonetheless, these results must be regarded carefully. This is because many tests of the TOH open the door for regression to the mean to generate spurious support (van Heerwaarden & Kellermann, 2020). Baseline thermal tolerance values are used to calculate plasticity, and then plasticity is modeled as a function of the baseline value. A common term is present on both sides of the function, potentially leading to a negative relationship that is a statistical artifact.

**Table 1.** Summary of the data sets reanalyzed in the present study. “Unadjusted” correlation coefficients were calculated using raw plasticity data as in the original analyses, and “Adjusted” correlation coefficients were calculated after removing the effect of regression to the mean (Kelly and Price 2005). Correlation coefficients (*r*) in bold indicate statistically significant relationships at the α = 0.05 level. If a study conducted multiple analyses with different subsets of their data, the subset is indicated by the “Sample” column. The “Metric” definitions are as follows: *CTmax* = critical thermal maximum; *LD50* = temperature of 50% mortality; *CCRT* = chill coma recovery time; *CCO* = chill coma onset.

| Clade | Species | Level | Metric | Sample | Unadjusted *r* | Adjusted *r* | Reference |
| --- | --- | --- | --- | --- | --- | --- | --- |
| **Heat tolerance** |  |  |  |  |  |  |  |
| Fish | *Gasterosteus aculeatus* | Individual | CTmax | - | **-0.59** | 0.02 | Mottola et al. 2022 |
| Fish | *Danio rerio* | Individual | CTmax | Females | **-0.70** | -0.20 | Morgan et al. 2018 |
| Fish | *D. rerio* | Individual | CTmax | Males | **-0.80** | 0.02 | Morgan et al. 2018 |
| Reptile | *Anolis sagrei* | Individual | CTmax | - | **-0.55** | 0.04 | Deery et al. 2021 |
| Reptile | *A. carolinensis* | Individual | CTmax | - | **-0.65** | 0.00 | Deery et al. 2021 |
| Reptile | *Lampropholis coggeri* | Individual | CTmax | Observer 1 | **-0.77** | **-0.49** | Phillips et al. 2016 |
| Reptile | *L. coggeri* | Individual | CTmax | Observer 2 | **-0.52** | -0.14 | Phillips et al. 2016 |
| Reptile | *Urosaurus ornatus* | Individual | CTmax | - | **-0.48** | -0.09 | Gilbert and Miles 2019 |
| Insect | *Drosophila melanogaster* | Population | CTmax | - | **-0.73** | **-0.65** | van Heerwaarden et al. 2016 |
| Crustacean | *Acartia tonsa* | Population | LD50 | - | **-0.84** | -0.57 | Sasaki and Dam 2019 |
| Crustacean | *A. tonsa* | Population | LD50 | - | **-0.97** | 0.66 | Sasaki and Dam 2020 |
| Crustacean | *A. tonsa* | Selection line | LD50 | CT pop., control | **-0.99** | **-0.97** | Sasaki and Dam 2021 |
| Crustacean | *A. tonsa* | Selection line | LD50 | CT pop., warming | **-0.94** | **-0.89** | Sasaki and Dam 2021 |
| Crustacean | *A. tonsa* | Selection line | LD50 | SP pop., control | **-1.00** | -0.93 | Sasaki and Dam 2021 |
| Crustacean | *A. tonsa* | Selection line | LD50 | SP pop., warming | **-0.99** | **-0.99** | Sasaki and Dam 2021 |
| Crustacean | *A. tonsa* | Selection line | LD50 | PG pop., control | **-0.97** | -0.91 | Sasaki and Dam 2021 |
| Crustacean | *A. tonsa* | Selection line | LD50 | PG pop., warming | **-0.98** | **-0.97** | Sasaki and Dam 2021 |
| **Cold tolerance** |  |  |  |  |  |  |  |
| Insect | *D. melanogaster* | Female lines | CCRT | Low heat | **-0.81** | 0.01 | Noh et al. 2017 |
| Insect | *D. melanogaster* | Female lines | CCRT | Mid heat | **-0.77** | -0.04 | Noh et al. 2017 |
| Insect | *D. melanogaster* | Female lines | CCRT | High heat | **-0.87** | 0.02 | Noh et al. 2017 |
| Insect | *D. melanogaster* | Individuals | CCRT | 15°C Acclimated | **-0.68** | -0.09 | O'Neill et al. 2021 |
| Insect | *D. melanogaster* | Individuals | CCRT | 25°C Acclimated | **-0.62** | -0.03 | O'Neill et al. 2021 |
| Insect | *D. melanogaster* | Individuals | CCO | 15°C Acclimated | **-0.63** | 0.00 | O'Neill et al. 2021 |
| Insect | *D. melanogaster* | Individuals | CCO | 25°C Acclimated | **-0.62** | 0.01 | O'Neill et al. 2021 |
| Insect | *Aphaenogaster picea* | Populations | CCRT | - | **-0.88** | 0.04 | Nguyen et al. 2019 |

Approaches for identifying, avoiding, and/or correcting for regression to the mean exist, but to date have rarely been applied to tests of the TOH. O’Neill, Davis, & MacMillan (2021) tentatively found support for the TOH among *Drosophila melanogaster* individuals with respect to cold tolerance, but from the correlation structure within their data, they inferred that the relationship was probably artifactual. Deery et al. (2021) found initial support for the TOH at the individual level among *Anolis* lizards with respect to heat hardening, but subsequently conducted a data resampling analysis (Ghalambor et al., 2015; Jackson & Somers, 1991) and concluded that the relationships were spurious. Finally, Gunderson & Revell (2022) used simulations to show how pervasive regression to the mean is when testing the TOH at macroevolutionary scales. They implemented a post-hoc method for adjusting plasticity data to account for regression to the mean (Kelly & Price, 2005), and found that doing so reduced support for the TOH in a previously-published dataset (Armstrong, Tanner, & Stillman, 2019; Gunderson & Revell, 2022). In sum, studies that attempt to account for regression to the mean in tests of the TOH have generally found that support for the hypothesis declines. How general is this outcome?

## Methods

I reanalyzed data from published studies that tested and found support for the TOH, focusing on work conducted at the intraspecific level (see Gunderson & Revell (2022) for analysis of the same issues at a macroevolutionary scale). I started with data from the studies summarized in Table 1 of van Heerwaarden & Kellermann (2020), and added as many subsequently-published data sets as I could identify. Data for some studies could not be obtained or the analysis supporting the TOH did not allow for correction. In total, I reanalyzed 25 correlations between baseline thermal tolerance and thermal tolerance plasticity representing 10 species (Table 1). I first conducted correlation analyses on the original data (“unadjusted”), and then with plasticity values adjusted to remove the effect of regression to the mean following the method developed by Kelly & Price (2005; “adjusted”). Detailed explanation of the method can be found in Kelly & Price (2005) and Gunderson & Revell (2022). All analyses were conducted in R (R Development Core Team, 2021). Data adjustment was done with a function I wrote in R (“rttm.adj”), and the script can be found in the Supplementary Materials.

## Results and Discussion

Two examples of what data look like before and after adjustment for regression to the mean are shown in Figure 1. The first example (Figure 1 A, B) shows heat hardening data for individuals of the lizard *Anolis carolinensis* (Deery et al., 2021). In the unadjusted data, there is a strong negative relationship between an individual’s baseline heat tolerance limit and how much their heat tolerance limit increases after heat shock (i.e., plasticity; Figure 1 A). However, that relationship disappears after plasticity values are adjusted for regression to the mean (Figure 1 B). The second example shows data on heat tolerance acclimation in *Drosophila melanogaster* (van Heerwaarden et al., 2016). In this case, a negative relationship between baseline tolerance and plasticity is maintained in both the unadjusted and adjusted data (Figure 1 C, D), though the effect of regression to the mean is still noticeable: the effect size (slope) and the correlation coefficient (and therefore variance explained; Table 1) are both slightly smaller using the adjusted versus the unadjusted plasticity data.

**Figure 1.**
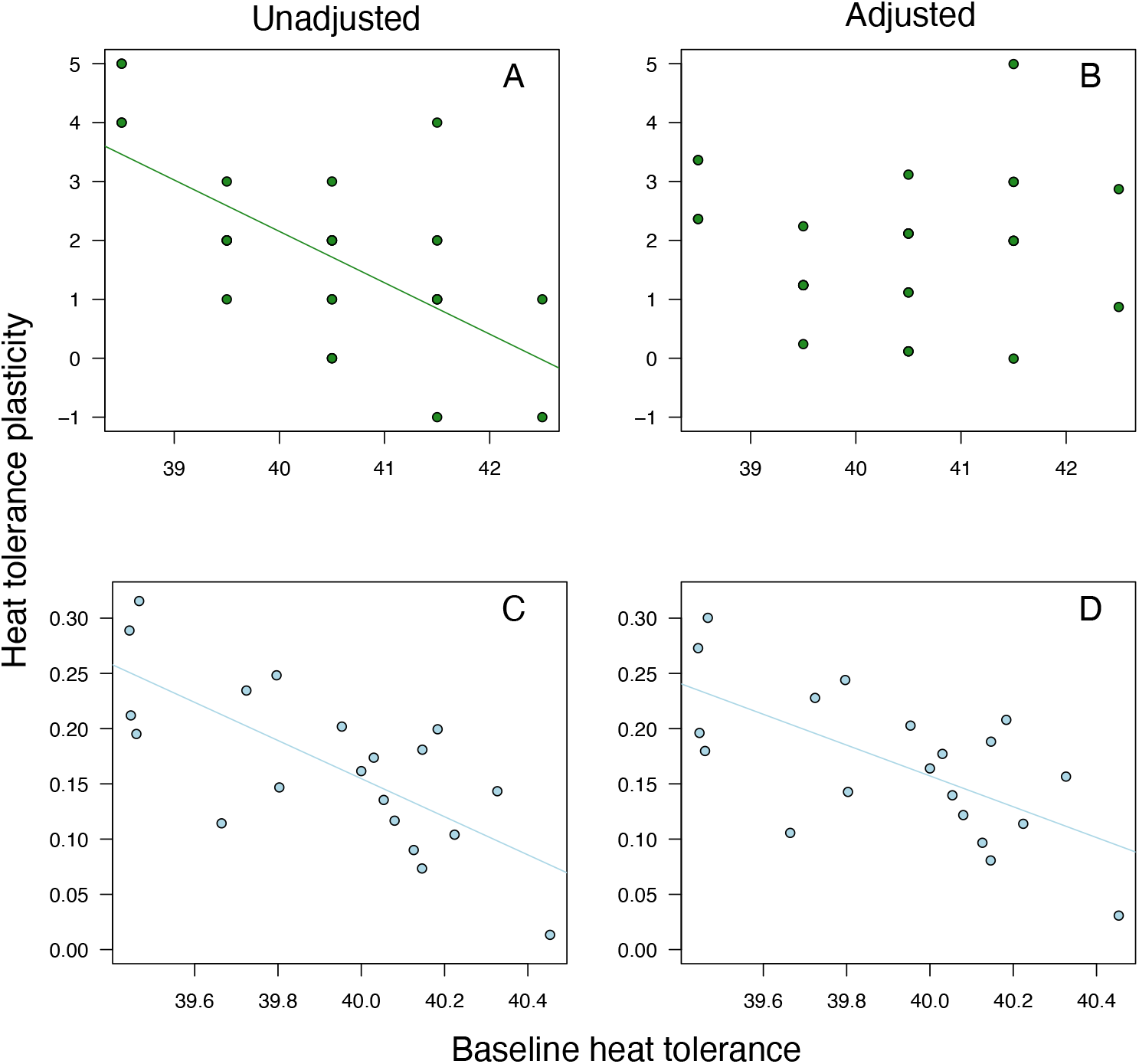
Examples of how tests of the Trade-off Hypothesis (TOH) are affected when plasticity values are adjusted to account for regression to the mean using the Kelly & Price (2005) method. A, B). Heat hardening data from the lizard *Anolis carolinensis* (Deery, Rej, Haro, & Gunderson, 2021). Support for the TOH was lost after data adjustment. C,D) Heat tolerance acclimation data from adult *Drosophila melanogaster* (van Heerwaarden, Kellermann, & Sgrò, 2016). Support for the TOH was maintained after data adjustment.

When this approach is applied across all the data sets, it becomes apparent that many relationships between baseline thermal tolerance and thermal tolerance plasticity are spurious due to regression to the mean (Table 1). Of the 25 results consistent with the TOH using unadjusted data, only six remain statistically significant at the α = 0.05 level after adjustment of plasticity values. Furthermore, correlation coefficients decrease in almost every case whether statistical significance is maintained or not. The magnitude and sometimes direction of effect also changes considerably after data adjustment. Slopes of the relationship between baseline tolerance and tolerance plasticity for all data sets become less negative when using adjusted versus unadjusted plasticity values (Figure 2).

**Figure 2.**
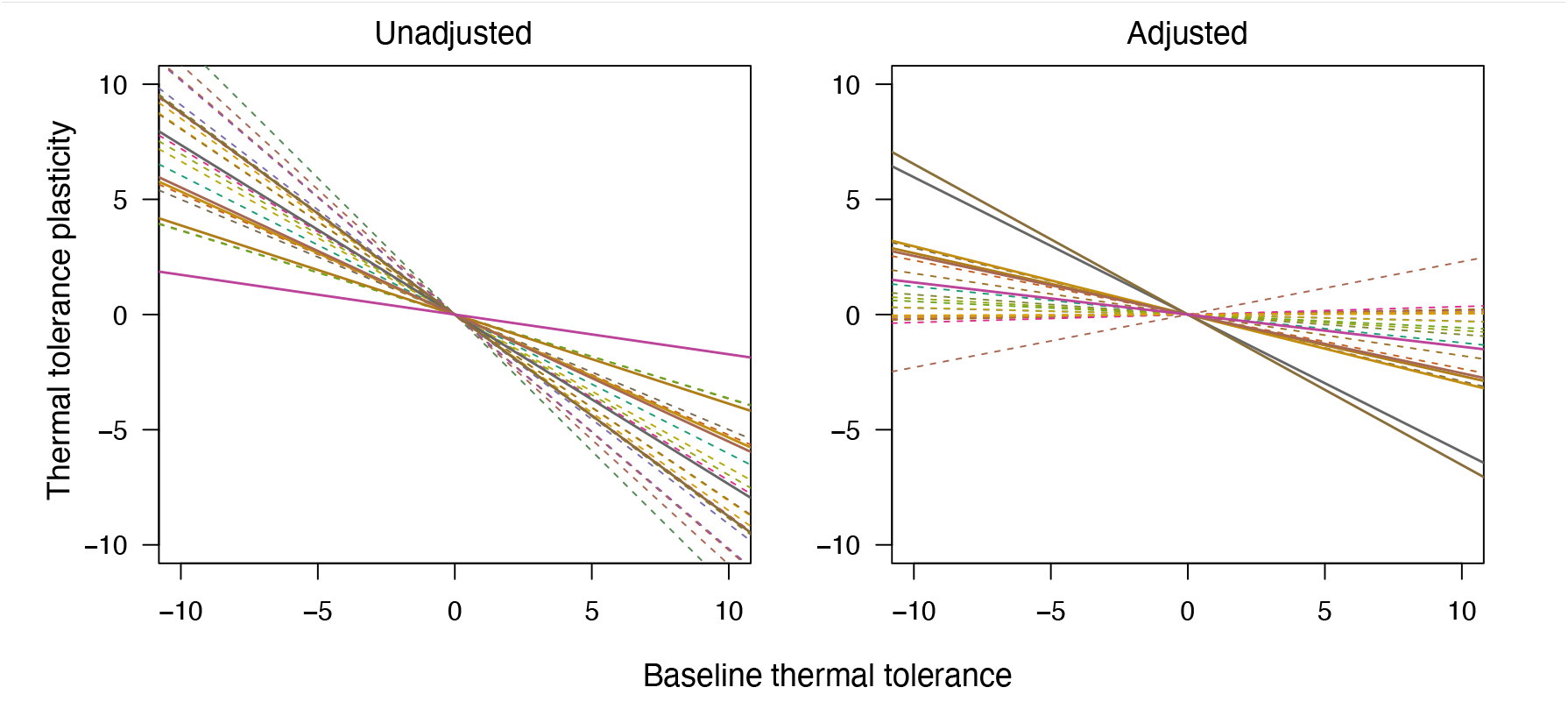
Slopes for the relationship between baseline thermal tolerance and thermal tolerance plasticity for the data sets in Table 1. Intercepts were set a 0 so all relationships could easily be visualized together. *Left*: Slopes when thermal tolerance plasticity values are not adjusted for regression to the mean. *Right*: Slopes when thermal tolerance plasticity values are adjusted for regression to the mean. Solid lines: data sets for which significance was maintained at α = 0.05 after data adjustment. Dashed lines: data sets for which statistical significance was not maintained after data adjustment.

Some of the regression to the mean effects demonstrated here are consistent with conclusions drawn by the original authors using different approaches (e.g., Deery et al., 2021; O’Neill et al., 2021; see above). However, most of the initial, unadjusted analyses were interpreted as evidence consistent with the TOH. Comparing data sets for which support for the TOH was and was not maintained after data adjustment is informative. Support for the TOH was rarely maintained after data adjustment in individual-level studies (Table 1). The lone example was with respect to heat hardening in the lizard *Lampropholis coggeri* (Phillips et al., 2016). Interpretation of this result is not straightforward, however, because it is not consistent with other data from the same species in the same study: two different observers measured thermal tolerance values, and support for the TOH was maintained after adjustment with data from one observer, but not the other (Table 1). Additionally, none of the analyses that remain statistically significant after data adjustment pertain to cold tolerance. Tolerance/plasticity trade-offs may be less common with respect to cold, but it is too early to draw broad conclusions because of the imbalance in effort devoted to the cold and warm ends of the spectrum. The TOH has been tested with respect to cold tolerance fewer times, in fewer species, and in fewer biological and experimental contexts (Table 1).

Data sets for which support for the TOH was maintained include a study of trade-offs between developmental and adult plasticity in *Drosophila melanogaster* (van Heerwaarden et al., 2016) and an experimental evolution study with the copepod *Acartia tonsa* (Sasaki & Dam, 2021). In the *Drosophila* study, animals were exposed to different developmental temperatures, after which these same animals had their capacity for adult thermal acclimation measured. The authors found that development in warm conditions led to increased baseline thermal tolerance, but lower capacity for adult acclimation (van Heerwaarden et al., 2016). A similar result was recently found with the frog *Lithobates sylvaticus*, in which acclimation to high temperature increased heat tolerance, but decreased capacity for rapid heat hardening (Dallas & Warne, 2023). Importantly, those authors used an experimental approach did not require post-hoc data adjustment (see Gunderson & Revell, 2022 for more details on experimental designs that avoid regression to the mean). The results of both studies are consistent with a mechanistic link between different forms of thermal plasticity (e.g., developmental plasticity, reversible acclimation, heat hardening) such that expression of one form reduces capacity to express another (Beaman, White, & Seebacher, 2016; van Heerwaarden et al., 2016).

In the experimental evolution study with the copepod *A. tonsa*, control and high temperature lines were established from three different source populations (Sasaki & Dam, 2021). In the initial analysis, there was an apparent trade-off between baseline thermal tolerance and plasticity over time across all control and high temperature groups from all three source populations. After data adjustment, the relationship remained statistically significant in all three of the high temperature selection groups, but in only one of the control groups (Table 1). Support for the TOH was also lost after adjusting data from two studies of thermal adaptation in natural *A. tonsa* populations (Sasaki & Dam, 2019; 2020; Table 1). That said, correlation coefficients remained relatively large and negative in all but one of the *A. tonsa* data sets in which statistical significance was not maintained (Table 1). In sum, there is compelling evidence for a real negative evolutionary correlation between baseline heat tolerance and heat tolerance plasticity in the *A. tonsa* system.

## Conclusion

Due to regression to the mean, support for the Trade-off Hypothesis is much lower than it appears at first glance (Table 1; see also Sgro et al. (2010) and van Heerwaarden, Lee, Overgaard, & Sgro (2014), who report analyses that reject the TOH with unadjusted data). Studies in which support for the TOH is maintained after data adjustment point to situations in which trade-offs between baseline tolerance and plasticity may be most common (Table 1; Dallas & Warne, 2023). However, at this point there are too few examples to make broad generalizations. More tests of the TOH are certainly warranted, but they must take regression to the mean into account using either post-hoc analyses or experimental designs that avoid the problem of regression to the mean altogether (discussed in Gunderson and Revell 2022). Of course, regression to the mean is not just an issue for tests of the TOH. It can also confound tests of other hypotheses in physiological ecology and evolution, including, but not limited to, those regarding plasticity in aerobic scope (Norin, Malte, & Clark, 2016) and trade-offs across time-dependent thermal tolerance landscapes (Jørgensen, Malte, & Overgaard, 2019). Researchers must be diligent.

At the core, inflated support for the TOH, or any other hypothesis, because of regression to the mean is a problem of identifying the appropriate null expectation (Hertz, Huey, & Stevenson, 1993). A lack of association between baseline thermal tolerance and thermal tolerance plasticity is not always the correct null hypothesis based on the methodological approaches taken (Gunderson & Revell, 2022). By recognizing such situations and accounting for them, we can more quickly develop a clear picture of the forces that influence the evolution of physiological plasticity, and in what situations different mechanisms are important. Doing so will greatly increase our understanding of organismal responses to ongoing anthropogenic warming.

## Acknowledgements

I would like to thank authors who provided data that were not publicly available. Eric Gangloff and two anonymous reviewers provided helpful feedback on the manuscript.

## Data availability

The data for this paper will be made available in Dryad.

