## Supplementary Material for "Trade-offs between baseline thermal tolerance and thermal tolerance plasticity are much less common than it appears"

**Supplementary Document**

R script for function to correct plasticity values for regression to the mean from Kelly and Price (2005). Takes two vectors, one for each phenotypic measurement used to calculate plasticity, and returns a data frame with three columns: *raw.plast* – unadjusted plasticity values. *dstar* – the correction factors to be applied to the mean raw plasticity value to create adjusted plasticity values. *adj.plast* – adjusted plasticity values.

```
rttm.adj<-function(m1, m2){
  raw.plast<-m2-m1
  vart<-var.test(m1,m2,paired = T)  ## variances equal?
  vpv<-vart$p.value  # var.test p value
  m1m2cor<-cor.test(m1, m2) # test correlation between m1 and m2
  rho<-m1m2cor$estimate # correlation coefficient between m1 and m2
  m1sd<-sd(m1) # m1 sd
  m2sd<-sd(m2) # m2 sd
  m1v<-var(m1) # m1 var
  m2v<-var(m2) # m2 var
  m1m<-mean(m1) # m1 mean
  m2m<-mean(m2) # m2 mean
  pm<-mean(raw.plast)
  rho2<-(2*rho*m1sd*m2sd)/(m1v+m2v) # adjusted correlation coefficient used if variances are equal
  rhof<-ifelse(vpv <= 0.05, rho, rho2) # which rho is used for dstar calculation is based on variance comparison
  dstar<-(rhof*(m1-m1m)-(m2-m2m))*-1 # adjustment values. Multiply by -1 to flip sign because Kelly and Price based on plasticity as m1-m2, not m2-m1 as in most thermal tolerance estimates
  adj.plast <- pm+dstar # corrected plasticity.
  out<-as.data.frame(cbind(raw.plast, dstar, adj.plast))
  return(out)
}
```

m1 = phenotypic measurement 1
m2 = phenotypic measurement 2

Kelly, C., & Price, T. D. (2005). Correcting for regression to the mean in behavior and ecology. *The American Naturalist, 166*(6), 700-707
